# Competition-by-drought interactions change phenotypic plasticity and the direction of selection on Arabidopsis traits

**DOI:** 10.1101/833848

**Authors:** Claire M. Lorts, Jesse R. Lasky

**Affiliations:** Department of Biology, Pennsylvania State University

**Keywords:** succession, tradeoffs, leaf economics, neighbor interactions, strigolactone

## Abstract

1. Populations often exhibit genetic diversity in traits involved in responses to abiotic stressors, but what maintains this diversity is unclear. *Arabidopsis thaliana* exhibits high within-population variation in drought response. One hypothesis is that competition, varying at small scales, promotes diversity in resource use strategies. However, little is known about natural variation in competition effects on Arabidopsis physiology.
2. We imposed drought and competition treatments on diverse genotypes. We measured resource economics traits, physiology, and fitness to characterize plasticity and selection in response to treatments.
3. Plastic responses to competition differed depending on moisture availability. We observed genotype-drought-competition interactions for relative fitness: competition had little effect on relative fitness under well-watered conditions, while under drought competition caused rank changes in fitness. Early flowering was always selected. Higher δ^13^C was selected only in the harshest treatment (drought and competition). Competitive context significantly changed the direction of selection on aboveground biomass and inflorescence height in well-watered environments.
4. Our results highlight how local biotic conditions modify abiotic selection, in some cases promoting diversity in abiotic stress response. The ability of populations to adapt to environmental change may thus depend on small-scale biotic heterogeneity.

## INTRODUCTION

The persistence of genetic variation in phenotypes in spite of the forces eliminating diversity is a central puzzle in biology (Mitchell-Olds *et al.*, 2007). While experiments often show strong selection under abiotic stress, populations still exhibit (perhaps surprising) diversity in response to such treatments (Hosgood & Parsons, 1968; Sultan & Bazzaz, 1993). Understanding diversity within populations is an important research goal partly because this diversity affects the adaptability of populations to environmental change (Pease *et al.*, 1989; Lynch & Lande, 1993; Jump *et al.*, 2009).

The effects of climate conditions (like precipitation) on organisms occur within the context of fine scale environmental heterogeneity, which can exacerbate or mitigate climatic conditions (Farrior *et al.*, 2013b; Urban *et al.*, 2013; Clark *et al.*, 2014; Moran & Alexander, 2014). This heterogeneity might select for phenotypic plasticity in a single optimal genotype. Alternatively, heterogeneity can promote genetic variation within populations by favoring different genotypes in different environments, *i.e.* a genotype-by-environment (G×E) interaction causing a fitness tradeoff across environmental gradients (Levene, 1953; Hamrick & Holden, 1979; Schmitt *et al.*, 2003).

Limiting resources that are influenced by climate, *e.g*. moisture, may be especially likely to exhibit interactions with competition. In plants for example, different levels of allocation to roots for water acquisition may be favored depending on the presence of neighboring individuals that compete for water (Farrior *et al.*, 2013a). Differing water use efficiencies (WUE) can be favored under moisture limitation depending on whether competitors are present (Campitelli *et al.*, 2016). Specifically, competition can favor more resource acquisitive strategies when resources are limited (Farrior *et al.*, 2013a).

Our current study system *Arabidopsis thaliana* (hereafter, Arabidopsis) is useful for studying these processes for several reasons. First, it exhibits substantial diversity in drought response within populations. Studies of genetic variation in δ^13^C, a proxy for water use efficiency (WUE), and stomata have found statistically significant but weak relationships with climate of origin (Dittberner *et al.*, 2018) as have studies of survival (Exposito-Alonso *et al.*, 2018). These weak relationships may be surprising given the tendency for local adaptation in Arabidopsis populations (Fournier-Level *et al.*, 2011) and stand in contrast with cold tolerance, which is more closely tied to local winter temperatures (Ågren & Schemske, 2012; Monroe *et al.*, 2016; Gienapp *et al.*, 2017). Second, while Arabidopsis occurs in a wide range of precipitation regimes (Hoffmann, 2002) it also occupies a wide range of microhabitats (Baron *et al.*, 2015), suggesting the opportunity for precipitation-by-microhabitat interactions. Some published data suggest the potential for competition-by-climate interaction effects on performance (Baron *et al.*, 2015; Campitelli *et al.*, 2016) though the physiological basis of these interactions is largely unknown. Third, previous research has yielded insight into genetic and physiological mechanisms of drought adaptation, providing a basis for specific hypotheses about how moisture and competitive gradients interact to select for different traits (McKay *et al.*, 2003; Juenger *et al.*, 2010; Des Marais *et al.*, 2012; Lovell *et al.*, 2013; Kenney *et al.*, 2014; Lasky *et al.*, 2014, 2018; El-Soda *et al.*, 2015; Bac-Molenaar *et al.*, 2016; Dittberner *et al.*, 2018; Exposito-Alonso *et al.*, 2018, 2019; DeLeo *et al.*, 2019).

Arabidopsis drought responses have been characterized into a stereotypical tradeoff between avoidance and escape (McKay *et al.* 2003; Kenney *et al.* 2014). Avoidance involves phenotypes that minimize water loss (*e.g*. stomatal closure) and maximize gain (*e.g*. root growth). Escape involves phenology and rapid development that matches with time periods when moisture is abundant (Ludlow, 1989). Escape may be adaptive when low soil moisture occurs at the end of growing seasons (Kenney *et al.*, 2014) while avoidance may be adaptive when moisture is limited throughout a growing season (Campitelli *et al.*, 2016). Escape has been characterized by rapid flowering, high specific leaf area (SLA), high leaf N, high carbon assimilation rate (*A*), low root:shoot mass ratio, and low water use efficiency (WUE) (Ludlow, 1989; Des Marais *et al.*, 2012; Kenney *et al.*, 2014; El-Soda *et al.*, 2015), while avoidance is characterized by the opposite values (but see Ferguson *et al.*, 2019). Arabidopsis is a rapidly maturing annual, an early successional strategy, which may be adaptive when neighbors are better resource competitors. In disturbed habitats where Arabidopsis frequently occurs, competition likely increases over time as colonists grow. In this situation, moisture availability may decrease across a season due to greater uptake combined with increasing evaporative demand as temperatures warm, favoring drought escape (Cohen, 1970; McLeod *et al.*, 2012; Campitelli *et al.*, 2016).

Despite the importance of Arabidopsis as the model plant, there exists little information on how diverse ecotypes respond physiologically to the presence of neighboring plants. Plant responses to neighbors may use a variety of cues, including nutrient availability (Schenk, 2006), red:far red light ratios (Aphalo *et al.*, 1999), or chemicals (Badri *et al.*, 2012). In Arabidopsis, Campitelli *et al.* (2016) found large biomass reductions and lower WUE in response to grass neighbors, even under ample water supply. One non-resource based mechanism for sensing neighbors is perception of root exudates, which are used by diverse species for belowground sensing of neighbors (Badri *et al.*, 2012; Schmid *et al.*, 2013; van Dam & Bouwmeester, 2016). Strigolactones (SLs) are exuded belowground by most plants (Al-Babili & Bouwmeester, 2015) and have been implicated in neighbor avoidance in a moss (Proust *et al.*, 2011). Furthermore, Arabidopsis plants can suppress inflorescence branching (Scaffidi *et al.*, 2014) and close stomata (Lv *et al.*, 2018) in response to SLs in growth media. To the extent that apical dominance in the presence of competitors enhances fitness (Baron *et al.*, 2015) it may be adaptive for a plant to suppress branching in response to SLs of neighbors.

We studied how competition interacts with moisture to drive plasticity and selection on life history and physiology in Arabidopsis. While additional resources vary with competition, *e.g.* nutrients, we controlled these in our experiments to focus on competition for water. We focused on these specific questions:

1. How does Arabidopsis respond to competition and drought via changes in life history and physiology? We hypothesized that drought would induce greater WUE due to stomatal closure, reduced performance, and greater SLA and leaf N as part of a drought escape strategy. We hypothesized that competition would induce inflorescence elongation, lower performance (biomass and fitness), and greater investment in resource acquisition (greater SLA, leaf N, lower WUE), partly due to perception of SLs in soil.
2. Does competition induce rank-changing G×E in fitness, suggesting that competitive gradients could promote diversity in response to abiotic stress within populations? We hypothesized that under conditions with ample resources, the presence of neighboring plants would have little influence on the relative fitness of diverse genotypes. By contrast, under moisture limitation we expected that competition would favor genotypes that differ in multiple traits compared to when competitors are absent.
3. What traits explain genetic variation in fitness response to competition (G×E)? We hypothesized that genetic tradeoffs are associated with alternate drought response strategies, such that contrasting treatments change the sign of directional selection coefficients across treatments.
4. Do traits exhibit genetic variation in plasticity that is under selection across competitive environments? We hypothesized that changes in some traits across competitive environments would be associated with higher average fitness across competitive environments.

## METHODS

### Plant material

We studied 17 diverse genotypes from across the native range of Arabidopsis and the reference genotype (Columbia) for a total of 18 genotypes (Tables S1 and S2). These genotypes mostly represent diverse lines used to create a multi-parent linkage mapping population (Kover *et al.*, 2009). We obtained seeds from the Arabidopsis Biological Resource Center (ABRC). However, it was later discovered that two lines (Can-0 CS76740, and Wu-0 CS78858) were contaminants from the supplying lab, *i.e.* they were not the original labeled genotype, though they were unique unidentified genotypes (Pisupati *et al.*, 2017) and we included them in the experiments. Seeds were grown for one generation in the lab at 16°C with a 16-hr photoperiod and the resulting next generation of seed was used in the present experiments. For competitors, we used three genotypes (Adi-10, Bd21-3, and Bd30-1) of *Brachypodium distachyon*, a widespread Mediterranean annual grass. Both Arabidopsis and *B. distachyon* occur broadly across mesic Mediterranean sites, making them potential natural competitors. However, these plant communities are diverse (Alados *et al.*, 2004) and it is likely selection on species interactions has been diffuse and not specific to this pair of species. The *B. distachyon* genotypes had similar growth form and rate and were simultaneously bulked in a greenhouse.

### Experimental design and conditions

To characterize genetic variation in response to moisture differences and their interactions with competition, we grew these genotypes in a split-plot design with two crossed treatments (for a total of four conditions): (1) well-watered, no competition (*i.e.* a single Arabidopsis plant in a pot), (2) well-watered, with competition, (3) intermittent drought, no competition, and (4) intermittent drought, with competition (Fig. S1). Each Arabidopsis genotype in each treatment had 10 total replicates per treatment, 5 for an initial harvest prior to flowering, and 5 for a post-senescence harvest (total Arabidopsis plants was 720).

We grew plants in a walk-in Conviron C1210 growth chamber (Conviron Ltd., Winnipeg, Canada). We designed a temperature and photoperiod program (Fig. S3, Table S3) to mimic a growing season starting with warm temperatures early in the local growing season, which we reasoned would be a potential site for moisture limitation, and centrally located in the species range (Online Supplement).

Competition was imposed by sowing six seeds of *B. distachyon*, along the outside perimeter of each pot. To focus on competition for water, we limited light competition by restraining *B. distachyon* tillers to the pot perimeter with a mounted stainless steel ring (Fig. S2). Drought treatments were imposed by removing bottom water at 16 days after planting (DAP), when all plants had at least the first true leaves fully expanded (Online Supplement).To limit nutrient competition pots in all treatments were fertigated at the same time of day with the same nutrition strength and volume.

#### Plant harvest and phenotyping during vegetative stage

To measure phenotypes during vegetative growth that required destructive harvest, we randomly selected five replicates for each genotype and treatment combination for harvest at 55-56 DAP, just prior to the initiation of the first flowering genotypes. In this harvest, *B. distachyon* shoots were cut at soil level, dried at 90 °C, and weighed. Roots of Arabidopsis plants under no competition treatments were washed, dried at 90 °C, and weighed (roots in competition were too difficult to separate from *B. distachyon* roots). Rosettes were removed from pots 1-2 cm below the soil level and were preserved in 75% ethanol at 4 °C. These rosettes were later measured for total leaf area, dry rosette biomass (excluding root), root diameter at 1 cm below the soil surface, stomatal density using a representative mid-aged leaf, specific leaf area (SLA) using one representative mid-aged leaf, leaf δ^13^C, leaf δ^15^N, and total leaf carbon and nitrogen composition. Leaf tissue samples for δ^13^C, δ^15^N, total carbon and nitrogen composition were prepared by drying the complete rosettes in coin envelopes for 12-24 hours at 65.6 °C, then grinding and homogenizing all leaves from the rosette to a fine powder using a mortar and pestle. Based on our estimated nitrogen content, we packaged a 1.5 – 2.0 mg aliquot of finely ground leaf tissue from each rosette into an 8 x 5 mm tin capsule (EA Consumables, Pennsauken, NJ). Samples were sent to the University of California Davis Stable Isotope Facility (SIF) for δ^13^C, δ^15^N, total carbon and nitrogen analyses using an elemental analyzer interfaced to a continuous flow isotope ratio mass spectrometer (IRMS). We measured whole-plant Arabidopsis gas exchange on a separate set of plants in an experiment that replicated the temperature, soil, and photoperiod conditions in the main experiment (Online Supplement).

#### Response to simulated root exudate strigolactones of neighbors

We used a synthetic strigolactone applied to the soil, GR24^rac^ (ChemPep, Inc., Wellington, FL), to explore how SLs from neighboring *B. distachyon,* which exude SLs from their roots (Changenet, 2018), might affect Arabidopsis performance and traits. Four of our genotypes were selected, two that were highly sensitive to competing grass neighbors (Sf-2 and Ct-1), and two that were less sensitive to neighbors (Wil-2 and Ws-0) under well-watered conditions. Plants were grown alone in pots (no competition). Growing conditions were as described in the main experiment above. Plants received bottom water until the first true leaves were expanded, at which point the water was removed and treatment applications were started. Plants were then top watered evenly throughout the top of the potting mix around the rosette using a pipette every day with 3 ml of either 7.8 uM GR24 solution, 0.78 uM GR24, or tap water, with 5 individuals per genotype per treatment. All plants were harvested 52 DAP, prior to flowering initiation. Rosette biomass and quantum yield of photosystem II using a MultispeQ V2.0 (Kuhlgert *et al.*, 2016) were measured. Leaf δ^13^C and total leaf C and N were measured from tissue analyzed at the UC Davis SIF.

#### Plant harvest and phenotyping at maturity

Fecundity of those Arabidopsis plants that survived to reproduction was estimated when all siliques reached maturity (dried and brown) and the rosette leaves had senesced, using total fruit length as an estimate of fecundity (Gnan *et al.*, 2014). We measured total silique number and length of six siliques corresponding to 10, 20, 40, 60, 80, and 90th percentiles of silique positions along the inflorescence in order to represent silique length along the inflorescence (*e.g*. 10th percentile silique was higher up the inflorescence than 10% of siliques). For each plant, we also measured inflorescence height and the number of inflorescence branches. After siliques were measured, we dried the mature inflorescence and rosette separately (in addition to rosettes that did not flower) at 90 °C for 24 hours to get aboveground dry biomass.

### Data analysis

Broad-sense heritability within treatments was calculated as the proportion of variation in phenotypes explained by genotype identity within each treatment, using a general linear model. We used linear mixed models to study sources of phenotype variation, with drought and competition treatments as fixed effects and genotypes, genotype-by-treatment interactions, and trays as random effects. With these mixed models we tested the hypothesis that fixed effects were zero and calculated variance components for random effects using the R package ‘VCA’ (Schuetzenmeister & Dufey, 2018). We calculated 95% confidence intervals on variance components (constrained to be non-negative) and report as significant components where the interval excludes zero. We calculated genotypes’ breeding values for each trait in each treatment using similarly constructed linear mixed models, using the R package ‘lme4’ (Bates *et al.*, 2013). To assess whether there were genetic correlations in traits often thought to represent coordinated strategies, we calculated Pearson’s correlations between breeding values for flowering time, SLA, leaf C:N, δ^13^C, and *A* (assimilation rate).

We tested for evidence of selection within each treatment by fitting linear regression models of relative fitness (absolute fitness / mean fitness among genotypes within a treatment) as a function of breeding values for individual unit-standardized phenotypes. Flowering time was a trait under strong selection in all treatments and thus we also tested trait correlations with fitness in multiple regression while also including a flowering time covariate, which allowed us to get better estimates of direct selection acting on a trait while accounting for flowering time selection (Lande & Arnold, 1983).

We tested for changing selection across competition treatments within the same moisture regime using t-tests. We focus on a soft selection scenario where individual fitness is density dependent, which may be particularly appropriate for comparisons across competitive treatments. Under this scenario, the appropriate approach is to relativize fitness within each treatment (*i.e.* combination of moisture and competitive conditions) (De Lisle & Svensson, 2017). For comparison, we included tests of changing selection when fitness was relativized across competitive treatments in the same moisture regimes, corresponding to a hard selection scenario (De Lisle & Svensson, 2017).

We also tested for selection on plasticity in response to competition, while accounting for differences among genotypes in their average traits. We fit linear models with breeding values and tested how trait in response to competition within a given moisture treatment was associated with absolute fitness in each individual competition treatment. We employed a multiple regression that included trait means averaged across competition treatments (Stinchcombe *et al.*, 2004). Greater detail on commands used in statistical software can be found in the Online Supplement.

## RESULTS

### Treatment effects on performance and physiology

We studied how drought and competition influenced individual Arabidopsis performance and physiology (Question 1 in Introduction). Drought and competition generally reduced plant performance, causing plants to have significantly smaller rosettes at mid-experiment harvest (rosette weight_vegetative_ and total leaf area_vegetative_), lower inflorescence weight, shorter inflorescences, fewer total mature fruits (siliques), shorter fruits, and less aboveground biomass (AGB_mature_ *i.e*. end of experiment harvest; linear mixed-effects models LMMs, all p < 0.05, Table 1, Figure 1). A notable exception was that plants had larger rosette weight_mature_ (at final harvest) under drought treatments compared to well-watered (LMM, drought fixed effect estimate = 55.3 mg, p < 10^-6^). The number of inflorescence branches was strongly reduced under competition (LMM fixed effect, p < 10^-8^), without any significant drought or interaction effects.

**Figure 1.**
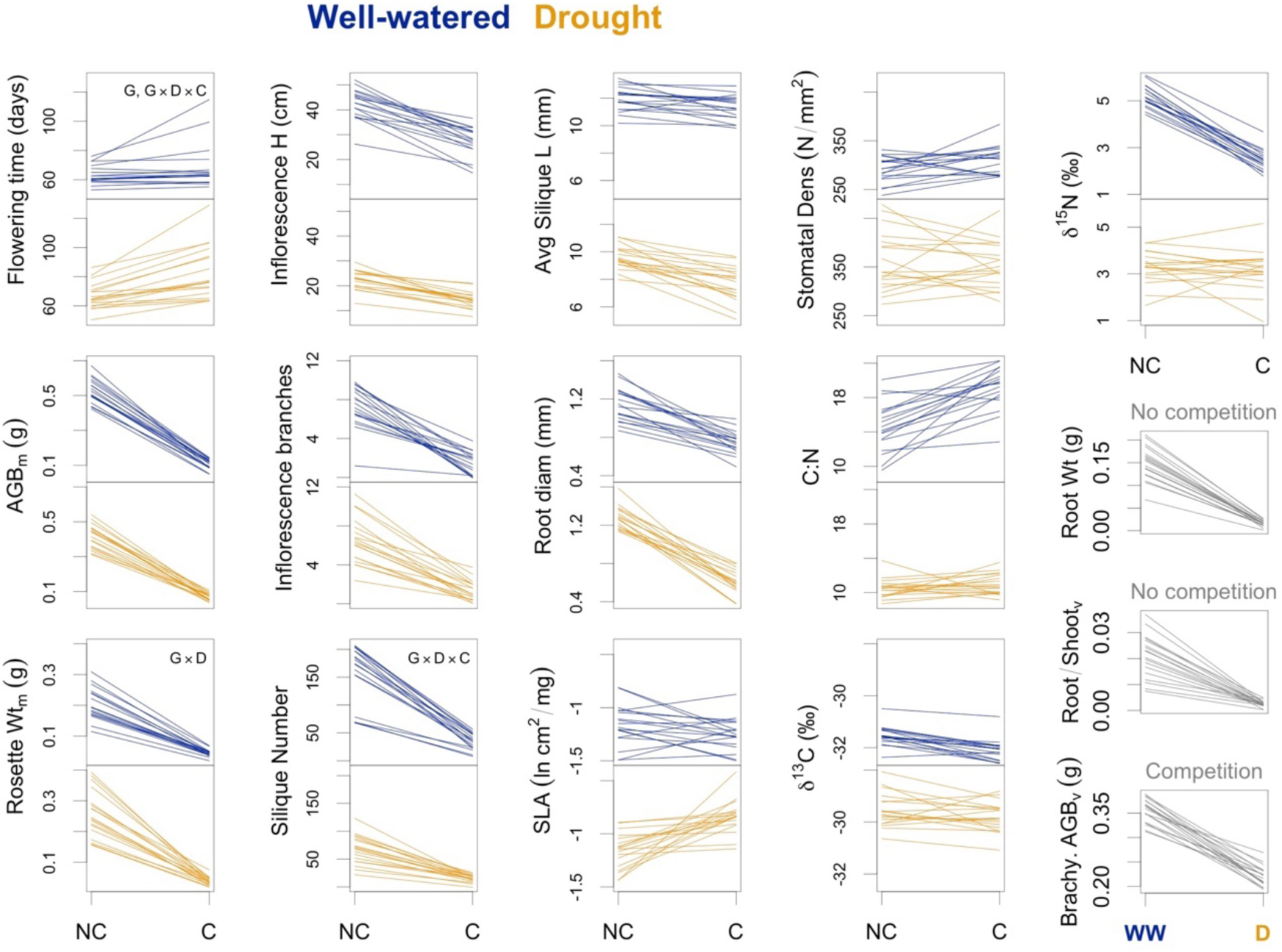
Reaction norms for traits across 18 Arabidopsis genotypes (each line represents one genotype). Lines show breeding values estimated from linear mixed models. ‘NC’ = no competition, ‘C’ = competition, ‘WW’ = well-watered, ‘D’ = drought. At right in gray are three traits that were only measured in a single competition/no-competition treatment. Three traits showing significant variance components for genotype (“G”) or genotype-treatment (e.g. “G×D”) interactions are indicated. All traits showed significant (FDR < 0.05) fixed effects for at least one treatment or treatment interactions (Table 1 for details).

**Table 1.**
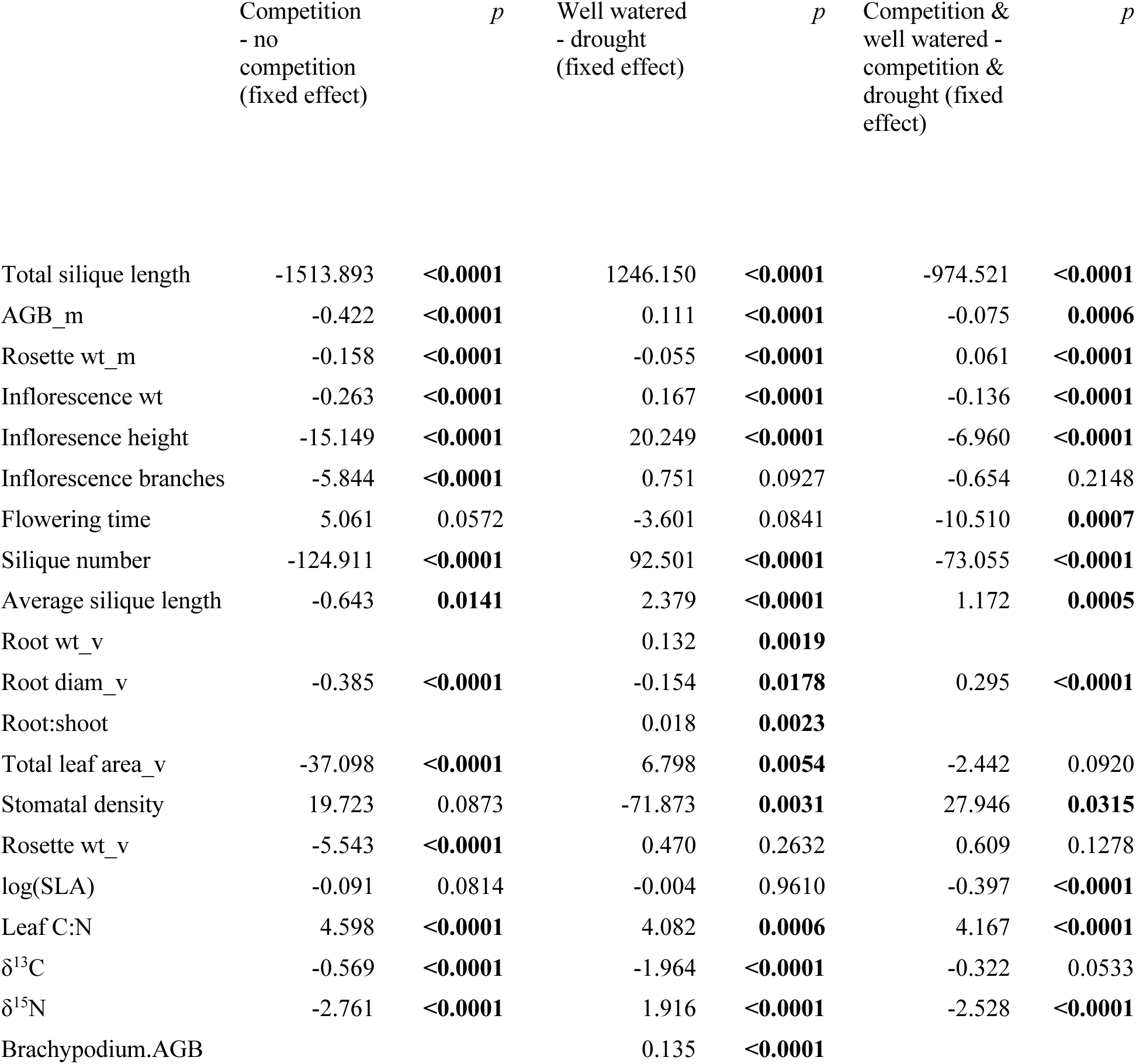
Results of treatment fixed effects on phenotypes from a linear mixed-effects model where genetic and tray effects were modeled as random effects. Fixed effects are in units of the trait. Bolded p values are significant at false discovery rate < 0.05 for each contrast across traits.

There was a significant interaction between competition and drought for AGB_mature_, inflorescence weight, inflorescence height, fruit number, average fruit length, such that under the combination of drought and competition these traits were greater than expected from the additive contribution of the two treatments (LMM fixed effects all p < 0.05, Table 1). However, rosette weight_mature_ and average fruit length were lower than expected under combined drought and competition (Figure 1, Table 1).

Survival to reproduction was high across treatments, with 95% of plants surviving until flowering (only one plant survived but did not flower by the end of the experiment). There was no pre-flowering mortality in either no-competition treatment (100% survival). By contrast, with competition, survival was 98% under well-watered conditions and 84% under drought, indicating that competition combined with drought was the harshest treatment. Mean genotype absolute fitness (mean estimated total mature fruit length per initial seedling following thinning) was highest under well-watered conditions without crowding (1900 mm, standard deviation among all individuals, SD = 599), followed by drought without crowding (656 mm, SD = 264), well-watered and crowding (387 mm, SD = 181), and drought and crowding combined (131 mm, SD = 95). We did not measure fecundity on *B. distachyon*, though its total aboveground biomass was greater in well-watered treatments compared to drought treatments at the mid-experiment harvest (mean pot total biomass, well-watered = 0.36 g, SD = 0.05; drought = 0.22 g, SD = 0.04).

In contrast to the whole plant performance measures described above, traits associated with life history or water balance physiology showed more diverse responses to water and competition (Figure 1). For example, mean flowering time showed a significant competition by drought interaction, such that flowering was significantly delayed only under combined competition and drought (LMM, competition×drought fixed effect p=0.0007, Table 1). Competition resulted in significantly decreased root diameter (LMM, fixed effect p < 10^-8^) and reduced δ^13^C (a proxy for water use efficiency, p < 10^-5^) across moisture treatments (Table 1). Under drought, competition resulted in even greater reductions in root diameter (LMM, competition×drought fixed effect p < 10^-6^) and greater SLA (p = 0.0002), while SLA was not affected by competition in well-watered conditions. Under well-watered conditions, competition resulted in greater leaf C:N (LMM, competition×drought fixed effect p < 10^-6^) and reduced δ^15^N (p < 10^-8^), while these were not affected by competition under drought (Table 1, Figure 1).

Drought treatments (apart from competition effects) resulted in reduced root weight, root:shoot ratio, greater stomatal density, lower leaf C:N, greater δ^13^C, and lower δ15N (all traits from mid-experiment vegetative harvest, LMM, all drought fixed effects p < 0.004, Table 1, Figure 1).

We also addressed whether plastic responses to competitor plants were consistent with effects of SLs in exudate. Consistent with responses to *B. distachyon* neighbors, Arabidopsis showed reductions in aboveground vegetative biomass with increased SL levels (GR24^rac^) under well-watered conditions (Figure S5, ANOVA on log proportional biomass with ranked GR24 treatment covariate, p < 10^-15^, genotype-GR24 interaction p = 0.7398). Howver, the SL treatment affected other traits in ways inconsistent with responses to *Brachypodium*, causing increased δ^13^C (ANOVA δ^13^C response to ranked GR24 treatment covariate, p = 0.0004, genotype-GR24 interaction p = 0.1588) and decreased leaf C:N (ANOVA, p < 10^-6^, genotype-GR24 interaction p = 0.2508, Figure S5). GR24 had genotype-specific effects on the quantum yield of photosystem II, with some genotypes having greater PSII yield with increasing GR24 and one having decreased PSII yield with increasing GR24 (ANOVA quantum yield of PSII response to ranked GR24 treatment covariate, p = 0.4047, genotype-GR24 interaction p = 0.0088).

### Genetic and genotype-environment interaction effects on traits

Broad-sense heritability within treatments differed widely among traits, with flowering time having the highest heritability in every treatment (ranging from 0.88 in drought and competition to 0.96 to well-watered and competition, Table S4). Heritability averaged among traits was highest in the well-watered no competition treatment, followed by well-watered competition, drought no competition, and drought competition, respectively. Heritability was higher for leaf C:N compared to leaf proportion N in most treatments and thus we below focus results on C:N.

We generally found that flowering time and δ^13^C were negatively correlated: later flowering genotypes had lower δ^13^C (well-watered, competition r = −0.50; drought, no competition r = −0.49, drought, competition r = −0.50) with the exception well-watered, no competition treatments where the two were unrelated (r= −0.07). As expected, SLA and leaf C:N were always negatively correlated (Figure S6). These traits were not closely related to *A* (assimilation rate) in well-watered conditions but under drought and no competition we observed that genotypes with greater *A* had greater SLA (r = 0.43), lower C:N (r = −0.56) and lower δ^13^C (r = −0.48).

We also tested for genetic variation in environmental responses to further address Question 1. Total fitness (Figure 2) and total silique number showed significant genotype-by-competition-by-water treatment interactions but not significant genotype interactions with competition or water alone (95% CIs of variance components excluded zero, Table S5), suggesting the potential for genetic tradeoffs in performance due to drought-competition interactions. Flowering time exhibited a strong genetic variance component and also a significant genotype-by-competition-by-treatment interaction but no genotype interaction with competition or water alone. Rosette weight_vegetative_ and rosette weight_mature_ both showed significant genotype-by-competition interactions but other traits did not (Figure 1, Table S5). Root:shoot, root weight, and δ^15^N (vegetative harvest) showed significant genotype variance components but no significant genotype-by-treatment interactions. Inflorescence height, inflorescence branch number, silique length, root diameter, SLA, stomatal density, C:N, δ^13^C, total rosette leaf area (vegetative harvest) did not show significant genotype-by-treatment variance components or significant genotype variance components (Table S5).

**Figure 2.**
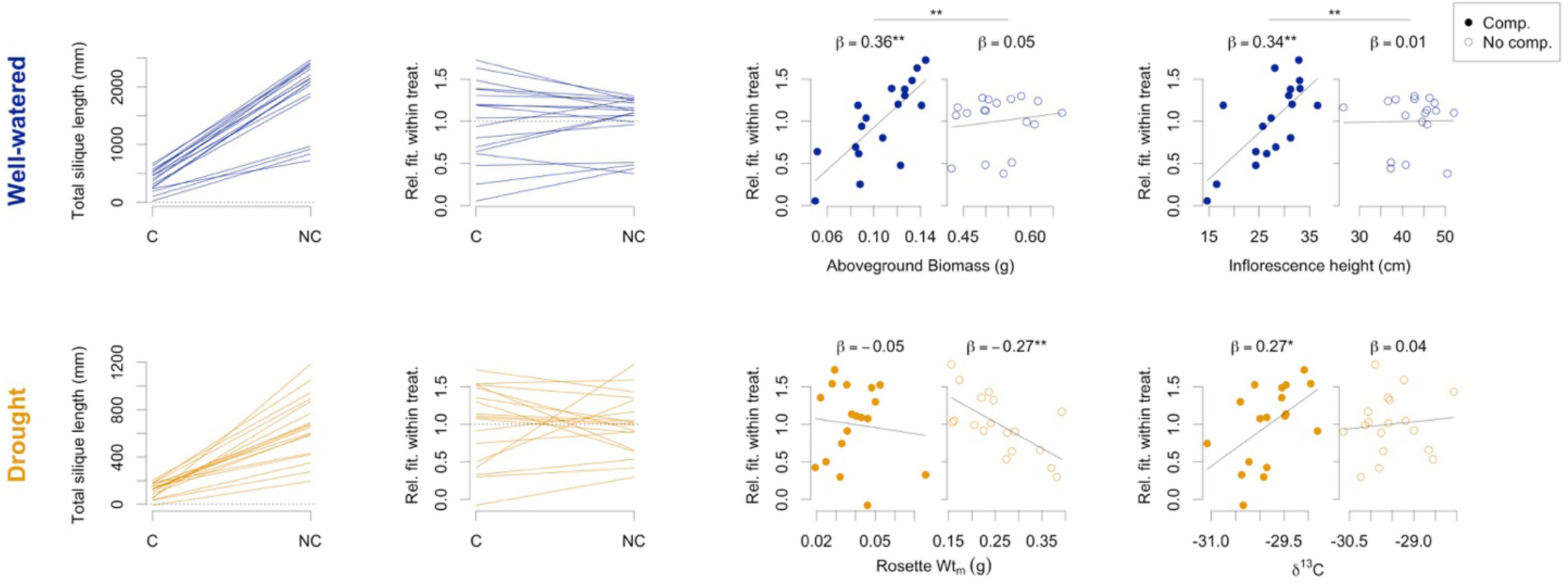
Reaction norms of absolute fitness (*i.e.* total silique length, first column) and relative fitness (standardized by dividing by mean fitness within each treatment, second columns) across competition treatments for the 18 studied genotypes (each line represents a genotype). Note that competition treatments are on the left and no competition on the right, so as to keep consistent x-axes for traits under selection (right columns). Lines show breeding values estimated from linear mixed models. The well-watered (top row) and drought (bottom row) treatments are shown (note their changing scales). Changing selection on traits among treatments (third and fourth columns). Standardized selection coefficients are shown, with * indicating nominal significance (p<0.05) and ** indicating FDR < 0.05. Horizontal bars with ** indicate slopes that differ significantly (t-test) between competition treatments in the same moisture regime with FDR < 0.05.

To assess the existence of fitness tradeoffs across environments (Question 2 in the Introduction), we calculated the correlation in relative fitness between competitive treatments. Under well-watered conditions, competition did not have a strong effect on the relative fitness of genotypes (absolute fitness correlation across environments, r = 0.82, p < 10^-5^, Figure 2). However, under drought, competition had a strong impact on relative fitness, such that absolute fitness was weakly related across competitive contexts (r = 0.38, p = 0.1170, Figure 2). Comparing moisture treatments under no competition, relative fitness was weakly correlated across environments, suggesting possible tradeoffs (r = 0.33, p = 0.1813). Under competition, fitness was more correlated across moisture treatments, indicating genotypes selected under competition have traits making them robust to moisture inputs (r = 0.52, p = 0.0263).

### Selection on phenotypes within and across environments

To identify traits affecting fitness variation (Question 3 in the Introduction), we tested for univariate and multivariate selection on traits within each environment. Earlier flowering time (FT) was favored in all four environments (linear models, well-watered, no competition p < 10^-6^, well-watered, competition p = 0.0020, drought, no competition p = 0.0162, drought, competition p = 0.0231, only well-watered significant at FDR < 0.05, Table 2). Greater inflorescence height (linear model p = 0.0001, significant after FDR) and inflorescence branch number (p = 0.0337, not significant after FDR) was linked to higher fitness in well-watered competition treatments but not in other treatments (all p > 0.05). Inflorescence height in well-watered competition treatments was selected even after conducting multiple regression also including a flowering time effect (inflorescence height p = 0.0214, flowering time p = 0.0469). Smaller rosettes (mature harvest) were selected under no competition treatments of both moisture regimes (well-watered, no competition p = 0.0095, drought, no competition p = 0.0030, both significant FDR < 0.05), though these patterns were not statistically significant after including flowering time in multiple regression (Table 2). AGB_mature_ was positively selected under well-watered competition treatments (p = 0.0002, significant after FDR), negatively selected under drought no competition (p = 0.0223, not significant after FDR), but unrelated with fitness in well-watered no competition treatments (p = 0.5625) and drought competition (p = 0.4926). Even after including flowering time in multiple regression, AGB_mature_ was positively selected in well-watered treatments with competition (AGB_mature_ p = 0.0020, FT p = 0.0159) but not so in drought treatments without competition (AGB_mature_ p = 0.4383, FT p = 0.2785). In the harshest environment, drought and competition, we found that δ^13^C was positively selected (p = 0.0291, not significant after FDR) though this was not significant after including FT (δ^13^C p = 0.1830, FT p = 0.1433). Stomatal density, SLA, root:shoot, root biomass, photosynthetic rate, C:N, δ^15^N, and root diameter did not show significant selection in any environment (Table 2).

**Table 2.**
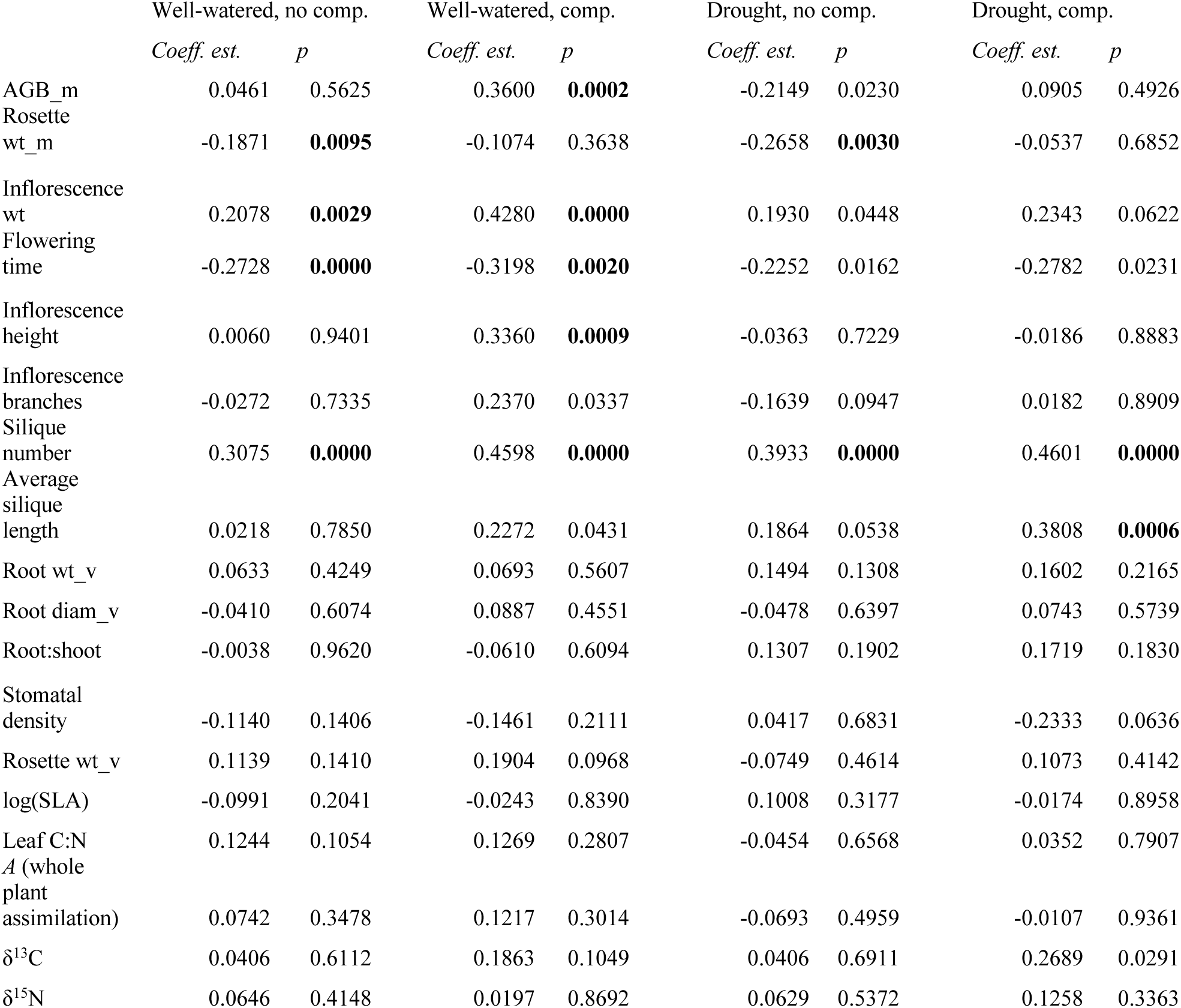
Standardized selection coefficients on breeding values for traits under different treatments. Bolded p-values are shown to indicate significance with false discovery rate < 0.05. Note that root weight, roo:shoot, and *A* were measured only on plants grown in the absence of competition. All other traits had breeding values and estimated fitness compared within each of the four treatments. All selection coefficients are in units of relative fitness (individual measures divided by the mean in each treatment) per standard deviation of the trait breeding values within treatment.

To determine the traits responsible for genotype-level tradeoffs across treatments (Question 3), we tested for changes in the coefficients of selection across competition treatments within a given moisture environment. When relativizing fitness within each treatment (De Lisle & Svensson, 2017), we found significant changes in selection depending on the competitive environment for inflorescence height and AGB_mature_ under well-watered conditions, with plants with taller inflorescences (two-tailed t-test p = 0.007, significant with FDR < 0.05), greater inflorescence mass (p < 0.0001, significant after FDR) and greater AGB_mature_ (p = 0.0003, significant after FDR) being more fit under competitive environments but with no detectable effect on fitness under no competition (Table 3, Figure 2). These changes in selection on these traits are surprising given the strong correlation (*i.e.* weak tradeoffs) between relative fitness of genotypes in well-watered conditions when comparing plants growing alone or with competitors (Figure 2). By contrast, we had observed changing ranks of relative fitness across competition treatments under drought (Figure 2). We did not find significant changes in selection on any phenotypes under drought across competition treatments (aside from the trivial case of silique number), though stomatal density showed the most substantial change (t-test p = 0.0630, Table 3). To further investigate the fitness tradeoffs across competition treatments under drought, we tested change in relative fitness between treatments and found it to be most closely associated with AGB_mature_ under drought and competition (relative fitness of drought under no competition - relative fitness under competition vs. AGB_mature_ under drought and competition, general linear model, p = 0.0118) while AGB_mature_ under drought no competition did not explain the fitness changes (p = 0.8333). When relativizing fitness across competition treatments in the same moisture environment (corresponding to a scenario of hard selection) (De Lisle & Svensson, 2017), flowering time in well-watered conditions was the only trait under significant changing selection across competition treatments, with relaxed selection under competition due to lower absolute fitness in this treatment (Table S6).

**Table 3.** Tests for changing selection between competition treatments within the same moisture treatment. Slopes give the relationship between traits and relative fitness, where relative fitness is standardized within the treatment combination of competition and moisture conditions (corresponding to a model of soft selection), and traits are standardized across competition treatments being compared. Bold indicates those where slopes differ significantly between competition treatments with FDR < 0.05. For results when fitness is relativized across environments see Table S6.

| *Trait* | *Well-watered* | *Slope, w/ comp* | *Slope, no comp.* | *t-test p* | *Drought* | *Slope, w/ comp* | *Slope, no comp.* | *t-test p* |
| --- | --- | --- | --- | --- | --- | --- | --- | --- |
| AGB_m |  | 2.77 | 0.15 | **0.0003** |  | 0.82 | -0.57 | 0.1886 |
| Rosette wt_m |  | -0.84 | -0.32 | 0.4907 |  | -0.50 | -0.43 | 0.9487 |
| Inflorescence wt |  | 2.40 | 0.47 | **<0.0001** |  | 1.58 | 0.47 | 0.1448 |
| Inflorescence height |  | 0.54 | 0.01 | **0.0070** |  | -0.03 | -0.05 | 0.9340 |
| Inflorescence branches |  | 0.69 | -0.04 | 0.0213 |  | 0.06 | -0.22 | 0.4723 |
| Flowering time |  | -0.25 | -0.52 | 0.0568 |  | -0.25 | -0.40 | 0.4410 |
| Silique number |  | 2.32 | 0.44 | **<0.0001** |  | 2.25 | 0.49 | **<0.0001** |
| Average silique length |  | 0.24 | 0.02 | 0.1244 |  | 0.43 | 0.31 | 0.5045 |
| Root diam_v |  | 0.19 | -0.06 | 0.3369 |  | 0.20 | -0.15 | 0.4690 |
| Stomatal density |  | -0.14 | -0.13 | 0.8890 |  | -0.27 | 0.04 | 0.0630 |
| Rosette wt_v |  | 1.15 | 0.16 | 0.0917 |  | 1.26 | -0.25 | 0.2883 |
| log(SLA) |  | -0.03 | -0.09 | 0.6743 |  | -0.03 | 0.14 | 0.4839 |
| A (whole plant assimilation) |  | 0.12 | 0.07 | 0.7313 |  | -0.01 | -0.07 | 0.7234 |
| Leaf C:N |  | 0.18 | 0.15 | 0.8567 |  | 0.03 | -0.05 | 0.6264 |
| $\delta^{13}\text{C}$ | | 0.22 | 0.05 | 0.3260 | | 0.29 | 0.04 | 0.1094 |
| $\delta^{15}\text{N}$ | | 0.07 | 0.20 | 0.7646 | | 0.11 | 0.07 | 0.7892 |

When testing for evidence of selection on traits averaged across competitive environments and independently on plasticity (Question 4 in Introduction), we found significant selection under well-watered conditions for a plastic increase in inflorescence height under competition compared to no competition (multiple linear regression, p = 0.0034) while average inflorescence height across treatments was not under significant selection, although this result was not significant after FDR control. For other traits where the average value was under selection across competitive environments, we found no independent selection on plasticity (Table S7).

## DISCUSSION

Organismal responses to abiotic conditions can be influenced by biotic interactions, such as interactions that affect general organismal health (*e.g*. parasitism, Price, 1991), change the balance of optimal allocation to abiotic stress tolerance (*e.g*. facilitation in arid systems, Bertness & Callaway, 1994), or directly change the availability of abiotic resources (*e.g*. competition for water, Cohen, 1970). Given rapid global climate change, we are particularly interested in limiting resources that are influenced by climate, like moisture. Such resources are likely to exhibit interactions between climate and competition in their effects on organisms. While a broad literature exists studying plant performance responses to drought combined with competition (e.g. Callaway & Walker, 1997; Young *et al.*, 2017), less is known about the underlying physiology and genetic variation. Here, we studied the model plant Arabidopsis to characterize plastic responses to drought and competition as well as to determine how competition can change selection for traits related to water balance. We found strong plastic responses to neighboring plants that differed between drought and well-watered conditions. Furthermore, we found that under drought there are major rank changes in relative fitness depending on the presence of competitors.

### Plastic responses to competition and moisture

We found plastic changes in most phenotypes, with strong interactions between competition and moisture. Competition and drought reduced performance in both well-watered and drought treatments, although plants in drought and competition had higher performance (biomass and total silique length) than expected from the additive contribution of each stressor. This interaction may indicate multiplicative growth responses to increased resources available in the absence of competition or with sufficient water. Additionally, plants under drought without competition generally had larger rosettes at maturity than in other conditions (Figure 1), perhaps signifying an adaptive conservative response that reduces reliance on inflorescence (cauline) leaves for photosynthesis.

By contrast, life history and physiology showed diverse plastic responses to our treatments. Generally, drought responses showed a strategy partly indicative of adaptive escape, *i.e.* a resource acquisitive strategy: lower leaf C:N and greater stomatal density, perhaps allowing plants to rapidly increase photosynthesis and take advantage of pulses of water. At the same time, higher δ^13^C under drought indicated higher water use efficiency and stomatal closure. Des Marais *et al.*, (2012) found that Arabidopsis tends to increase SLA and decrease C:N under drought, which the authors interpreted as drought escape. However, several non-vernalization requiring genotypes in Des Marais *et al.* (2012) showed opposite patterns: greater C:N and lower SLA under drought. Our study genotypes were non-vernalization requiring, and we found that drought increased SLA only under competition. There was no drought effect on SLA when plants were alone. These drought-by-competition interactions suggest the escape strategy might be specific to certain life history strategies or combined competitive and drought conditions.

Responses to competition differed between drought and well-watered conditions, demonstrating the interdependence of abiotic and biotic effects. Under well-watered conditions, response to competition was characterized by higher leaf C:N, suggesting a decrease in a photosynthetic rate per leaf mass, although this was coupled with a consistent but slight decrease in δ^13^C, the latter of which suggests an acquisitive growth strategy via greater stomatal or mesophyll conductance to escape competition and is consistent with findings by Campitelli *et al.* (2016). By contrast, competition under drought increased SLA, with some genotypes also reducing δ^13^C, perhaps indicating an acquisitive escape strategy (allowing greater whole-plant carbon assimilation) (Des Marais *et al.*, 2012). We found that under drought, genetic variation in SLA was positively associated with whole plant carbon assimilation (Figure S6). In one of the few relevant studies, Ehleringer (1993) found that the desert shrub *Encelia farinose* (Asteraceae) showed no δ^13^C response to competitor removal.

δ^15^N was elevated in well-watered no-competition treatments, which we interpreted as a shift in Arabidopsis use of N pools among treatments. Synthetic fertilizers (*e.g.* Miracle Grow used here) cause lower leaf δ^15^N relative to organic sources like peat humus (used in our media), as fertilizer has more ammonium, which requires a single assimilation event (in roots) versus multiple assimilation events (in roots and shoots) for soil nitrate sources (Evans *et al.*, 1996; Bateman *et al.*, 2005; Trandel *et al.*, 2018). We infer that well-watered Arabidopsis plants without competition used more N from the peat humus versus fertilizer. Under well-watered conditions with competitors, the fertilizer N source may have been favored by the Arabidopsis plants due to smaller root systems for exploitation of soil N, causing low leaf δ^15^N. Under drought, N uptake may have been restricted to brief periods following fertigation, favoring the fertilizer source and causing low leaf δ^15^N.

Even under well-watered conditions with fertilizer and direct light, Arabidopsis responded to competition with reduced performance. Such a strong response suggests Arabidopsis plants were using a chemical cue, such as root exudates (Pierik *et al.*, 2013), to sense the neighboring *Brachypodium* (Badri *et al.*, 2012; Schmid *et al.*, 2013). Previous studies have shown that SLs in growth media can suppress inflorescence branching by Arabidopsis (Scaffidi *et al.*, 2014) and cause stomatal closure (Lv *et al.*, 2018). Our treatments with a synthetic SL reduced vegetative rosette biomass, suggesting SL signaling could partly explain Arabidopsis response to neighbors. The yield of PSII was greater for two genotypes under SL treatment, consistent with findings in tomato that the synthetic SL GR24 up-regulated light-harvesting genes (Mayzlish-Gati *et al.*, 2010). However, two other genotypes showed *lower* PSII yield in response to SL; these genotypes had lower biomass response to SL, for reasons that are unclear. We found that experimental SL addition caused an increase in δ^13^C (likely signaling stomatal closure, Lv *et al.*, 2018) and a decrease in C:N, patterns that were opposite the Arabidopsis response to *Brachypodium* in our well-watered treatment. This result indicates that other non-SL cues are required to explain the responses to *Brachypodium* that we observed under putative non-resource limited conditions (our well-watered treatment).

### Genetic correlations among traits

We found that genetic variation in flowering time was negatively associated with leaf δ^13^C, suggesting that early flowering plants had higher intrinsic leaf-level WUE due to lower stomatal conductance or higher assimilation rate *A* (Ferguson *et al.*, 2019). This positive correlation between flowering time and δ^13^C might be a result of maladapted late-flowering genotypes having lower *A* for a given stomatal conductance. Our findings contrast with some previous studies that found the early flowering plants had low leaf δ^13^C (McKay *et al.*, 2003; Lovell *et al.*, 2013; Kenney *et al.*, 2014). Our panel of natural genotypes lacked genotypes requiring vernalization, thus these genotypes might partly be responsible for the discrepancies with these publications. Additionally, Ferguson *et al.* (2019) did not find a flowering time relationship with leaf δ^13^C in comparisons of NILs at major flowering time QTL.

### Selection under competition and changes in selection depending on competition

Researchers have more commonly investigated how competition-drought interactions affect different species in a community (Marks & Strain, 1989; Miranda-Apodaca *et al.*, 2015; Napier *et al.*, 2016), or have studied changing selection driven by the interaction between drought and herbivory (Sthultz *et al.*, 2009; Huttunen *et al.*, 2010; Grinnan *et al.*, 2013) or competition and herbivory (Tiffin, 2002). However, there are few studies of changing selection due to drought-competition interactions. In one of the few such studies, Ehleringer (1993) found that in the desert shrub *Encelia farinose*, δ^13^C was not under selection in the presence of competitors.

However, upon competitor removal, lower δ^13^C was associated with higher growth (but not survival) possibly due to increased soil moisture. Here, we observed significant genotype-by-competition-by-moisture treatment interactions for performance related traits (fitness components and aboveground biomass) and flowering time. The genotype-by-competition-by-moisture effects on fitness indicate that the effects of competition on relative fitness are conditional on moisture environment. This competition-by-moisture effect on selection could be seen in the substantial changes in relative fitness depending on competition under drought, but the fairly consistent relative fitness despite the presence of neighbors in well-watered conditions (Figure 2). Rank changes in relative fitness across treatments differing in competition were observed among natural Arabidopsis genotypes at a moist field site (Baron *et al.*, 2015) and among genotypes segregating for a single water-use-efficiency locus under greenhouse drought (Campitelli *et al.*, 2016). Here we show selection among natural genotypes under drought. In nature, the heterogeneity of competition may select for alternate Arabidopsis traits in water limited environments, suggesting a potential mechanism promoting their high genetic diversity in drought responses (Dittberner *et al.*, 2018; Exposito-Alonso *et al.*, 2018).

We identified several traits associated with fitness in specific competitive contexts. Inflorescence height and weight as well as AGB_mature_ showed the strongest evidence for changing selection depending on competition, both when well-watered. Under well-watered competition, plants with greater AGB_mature_ and larger inflorescences had greater fitness, but not in the absence of competition. Selection for AGB_mature_ may indicate that the investment in leaves for resource capture (comprising much of non-reproductive of AGB, and including both rosette and cauline leaves) is adaptive when surrounded by competitors, while allocation to reproduction may be selected when competitors are absent. The lack of selection for greater AGB_mature_ and taller inflorescences in drought may indicate that aboveground investments are less useful when competitors are smaller. We also found positive selection for the plastic increase in inflorescence height under well-watered competition treatments. Thus, inflorescence height may only be a good measure of fecundity under the treatment with the most competitor biomass (well-watered), where competitors lead to suppressed inflorescence branching (Schmitt *et al.*, 2003; von Wettberg *et al.*, 2008).

Under drought and competition, plants with greater δ^13^C had higher fitness (though this was not significant after FDR control), indicating greater leaf-level water use efficiency was adaptive, but not in other treatments. We expected lower WUE would be favored under drought with competitors to rapidly complete development before competitors depleted resources too low (Campitelli *et al.*, 2016). Instead, higher WUE may have been favored because conditions were too dry to permit success of a rapidly developing drought escape strategy, thus higher WUE may be necessary in these environments. Alternatively, despite the dramatically lower performance of plants in response to competition under drought, the intermittent watering may have been enough to favor conservative drought avoidance strategies (high δ^13^C). Our findings contrast with those of Kenney *et al.* (2014) who found terminal drought and well-watered treatments selected for lower δ^13^C and plants with greater AGB_mature_, suggesting escape strategies were favored. Here, early flowering was favored in competitive treatments as it was in all treatments, similar to the consistently observed selection for earlier flowering in many experiments (Austen *et al.*, 2017) including those on Arabidopsis under terminal drought (Kenney *et al.*, 2014).

### Conclusions

The effects of abiotic stress on phenotypic plasticity and natural selection depend on biotic context. Arabidopsis represents an important model system for plants, including for its responses to environment. Our results highlight how natural variation in environmental responses should be interpreted in light of biotic interactions.

## Acknowledgements

Jonathan Kizer, Sarah Lucas, and Crosley Kudla-Williams assisted in harvesting and phenotyping. David Des Marais provided initial seeds of *Brachypodium distachyon*.

## Author contributions

CML carried out experiments, analyzed data, and wrote the manuscript. JRL planned the experiments, analyzed data, and wrote the manuscript.

## SUPPLEMENTAL TEXT, FIGURES, AND TABLES

### Growth conditions

Specifically, we chose conditions for the site of collection of the Lip-0 ecotype near Chrzanów in southern Poland. We used a growing season model (Lasky *et al.*, 2012) with WorldClim data (Hijmans *et al.*, 2005) then extracted a monthly time series of maximum temperatures and minimum temperatures. We used these temperatures to program the growth chamber, though in the early growing season we were limited to 10 °C nightly temperature because the chamber was unable to generate temperatures < 10 °C. Local photoperiod for the growing season was obtained using the ‘maptools’ package (Bivand & Lewin-Koh, 2019). Weekly conditions were obtained via linear interpolation of the monthly temperature values (Table S2). Our growth chamber had constant horizontal airflow to maintain homogenous conditions throughout.

Plants were grown in SC-10 Cone-tainers^TM^ (3.81 cm diameter, 164 ml) in a 98-cell rack (RL98) leaving adjoining cells in the tray empty to prevent shading interaction between neighboring pots (Stuewe and Sons, Tangent, Oregon, USA). Each rack was placed within individual flow trays. Competition treatments were spaced at every other pot location to prevent among-pot crowding, and each tray was designated a drought or well-watered treatment (Fig. S1). Genotype pot locations were randomly assigned. Potting composition within each pot consisted of 50% potting mix (Premier pro-mix PGX), 25% medium commercial grade sand, 25% calcined clay (TURFACE® ALL SPORT PRO™, Turface Athletics™, PROFILE Products LLC, Buffalo Grove, IL), with 2.5 cm of the potting mix as a topsoil to assist Arabidopsis plants in germination and initial growth. Glad Press’n Seal Wrap (The Glad Products Company; Oakland, CA) was placed over pots until germination to prevent seed dehydration prior to germination.

Germination for *B. distachyon* was near 100%. The three *B. distachyon* genotypes were randomly sorted into competition pots. Preliminary experiments using varying levels of *B. distachyon* in competition with *Arabidopsis* showed that greater than six plants resulted in overcrowding and very poor growth of *B. distachyon,* thus six plants were used as a competition treatment in pots.

Pots in the drought treatment were not watered during 17-19 DAP prior to fertigation (combined watering and fertilizing) treatments to allow for initial drying. To simulate persistent rainfall limitation, starting at 20 DAP, each individual pot in the drought treatment was fertigated from above with 3 ml of full strength Miracle-Gro solution (Miracle-Gro® Water Soluble All Purpose Plant Food, The Scotts Company, LLC., Marysville, OH) every other day. Throughout the entire experiment, all pots in all treatments were fertigated at the same time of day with the same nutrition strength and volume to limit nutrient competition and to focus on traits involved in competition for water. At 29 DAP, fertigation was shifted to 6 ml of ½ strength Miracle Gro solution, and at 50 DAP fertigation was increased to 12 ml of ½ strength Miracle Gro solution. Well-watered pots were kept consistently moist by bottom watering with tap water. We defined the end of the plant life cycle as the time when 50% of siliques were mature (brown and brittle), because at this stage the ecotypes used in this experiment stopped producing new flowers and discontinued development of existing fruits if present. Bottom water in well-watered treatments was removed when at least one plant had 50% mature (brown and brittle) siliques, and the remaining flowering plants were top watered daily with 12 ml of ½ strength Miracle Gro, to maintain the potting mix water and nutrient content of the well-watered plants, until 50% of siliques were mature. Watering in drought treatments was also discontinued for a pot when 50% of siliques were mature. Potting mix volumetric water content for each treatment was monitored throughout the entire experiment with EM-50 dataloggers and EC-5 probes (METER Group Inc., Pullman, WA) within designated non-experimental pots with the reference genotype Col-0 (Fig. S4).

### Photosynthesis

We measured whole-plant gas exchange on a separate set of plants in an experiment that replicated the temperature, soil, and photoperiod conditions in the main experiment. However, we did not include *B. distachyon* competitors and we covered topsoil in clay. We did not use competition treatments due to the challenges of measuring whole plant gas exchange on the focal Arabidopsis plant when surrounded by *B. distachyon* competitors. We used the clay cap as suggested by the gas exchange analyzer manufacturer (LI-COR Inc., Lincoln, NE, USA) to limit soil respiration from interfering with photosynthetic gas exchange measurements. Prior to planting, the top soil was covered with a 0.5 cm layer of potting clay (Msanne *et al.*, 2011; Souza *et al.*, 2015) with a 0.5 cm hole in the center for plant growth. Even with the clay and initial soil sterilization, we observed low, non-zero soil respiration from well-watered pots, and as a result below we do not focus analysis on changes in photosynthesis between treatments, but instead use photosynthesis measurements in selection analyses within moisture treatments.

Five replicates per genotype per water treatment were planted in a randomized block design. Photosynthesis (*A*) was measured between 8:00 AM and 12:30 PM using a LiCor 6400XT with a LI6400-17 Whole Arabidopsis Chamber (LI-COR Inc., Lincoln, NE, USA) modified to fit SC-10 Ray Leach Cone-tainers™ (Stuewe and Sons Inc., Tangent, OR) by widening the opening that holds the Cone-tainer to fit a size -223 O-ring (McMaster Carr part number 1170N95, Elmhurst, IL). This modification created an air-tight seal around the circumference of the Cone-tainer approximately 4 mm from the top of the Cone-tainer, to prevent gas leaks from the surrounding environment and to allow for continuous air flow and even concentrations of gasses throughout the chamber during measurements.

### Statistical analysis

We used linear mixed models to test fixed effects of treatments on Arabidopsis traits, as well as to test for significance of variance components for genotype and genotype-treatment interaction random effects. To do so, we used the VCA package for R. Example code is shown below, where ‘mydata’ is a data.frame with rows as individual plants and columns that include a phenotype variable, ‘CompTRT’ and ‘WaterTRT’ variables that are factors giving the competition and moisture conditions respectively, and ‘Tray’ giving the identity of the tray the plant was in.

~~~
library(VCA)

model1 <- anovaMM(phenotype ∼ CompTRT * WaterTRT * (genotype) +(Tray),
   Data = mydata)

inf <- VCAinference(model1, VarVC=TRUE)

VCAinference(model1)

lc.mat <- VCA::getL(model1, c("CompTRTC-CompTRTNC", "WaterTRTWW-
   WaterTRTD", ’CompTRTC:WaterTRTWW-CompTRTC:WaterTRTD’))

test.fixef(m1a, lc.mat)
~~~

We used linear models to test for selection on traits within each treatment. Example code is shown below, where ‘mydata2’ is a data.frame with rows as genotypes, ‘phen’ are breeding values for a given phenotype in a given treatment, and ‘RelFit’ is fitness relativized to the mean within that treatment:

~~~
model2 <- lm(RelFit ∼ phen, data = mydata2)
summary(model2)
~~~

We also used linear models with t-tests for changes in selection among competition treatments, within the same moisture treatment. Example code is shown below, where ‘mydata3’ is a data.frame with rows as genotypes, each genotype having two rows, one for breeding values without competition in a given moisture environment and one for breeding values with competition in a given moisture environment. ‘phen’ is a column giving breeding values, ‘compT’ is a factor giving the competition treatment, and ‘RelFit’ is fitness relativized to the mean within that treatment:

~~~
model3 <- lm(RelFit ∼ phen * compT, data = mydata3)
summary(model3)
~~~

We used linear models again to test for selection on competition plasticity while accounting for average trait values. Example code is shown below, where ‘mydata3’ is a data.frame with rows as genotypes. ‘avg_phen’ is a column giving breeding values averaged across competition treatments for a given moisture environment, ‘plast_phen’ is a column giving the difference in breeding values between competition treatments for a given moisture environment, and ‘RelFit’ is fitness averaged across competition treatments for a given moisture environment and then relativized to the mean.

~~~
model4 <- lm(RelFit ∼ avg_phen + plast_phen, data = mydata4)
summary(model4)
~~~

**Figure S1.**
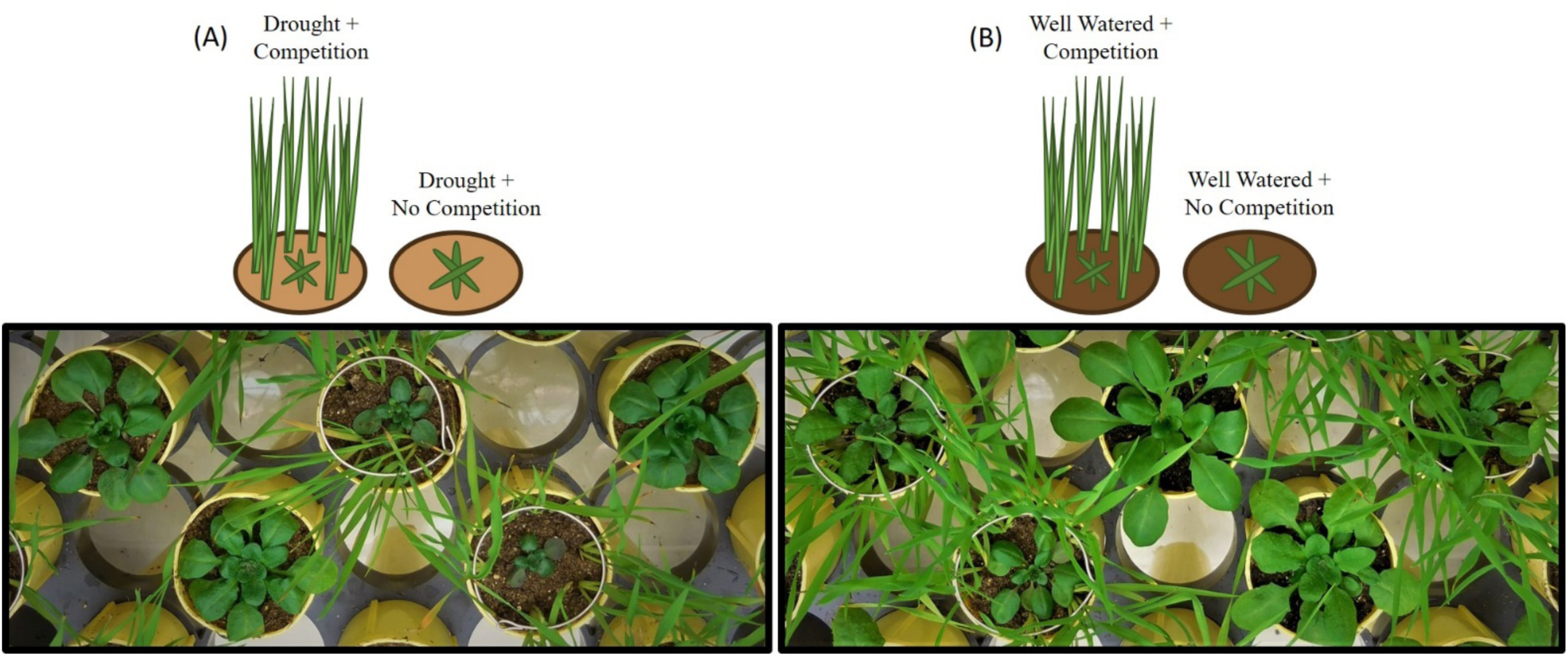
Experimental treatments, including (A) drought treatment under competition and no competition, and (B) well-watered treatment under competition and no competition. Photographs were taken 53 days after planting, prior to the first harvest.

**Figure S2.**
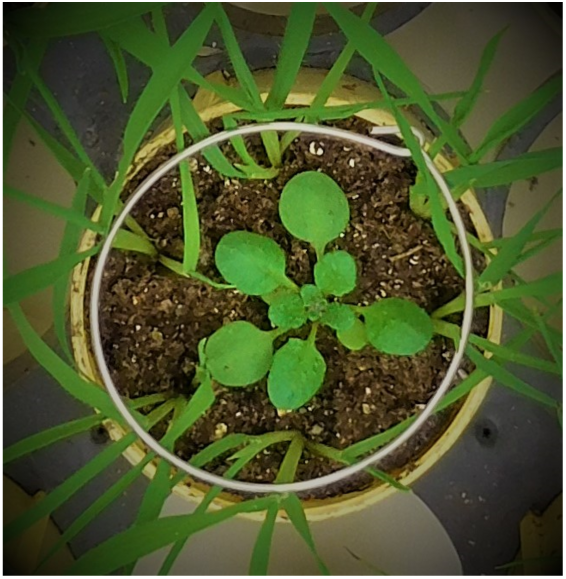
Experimental plant with stainless steel wire installed to prevent *B. distachyon* shading.

**Figure S3.**
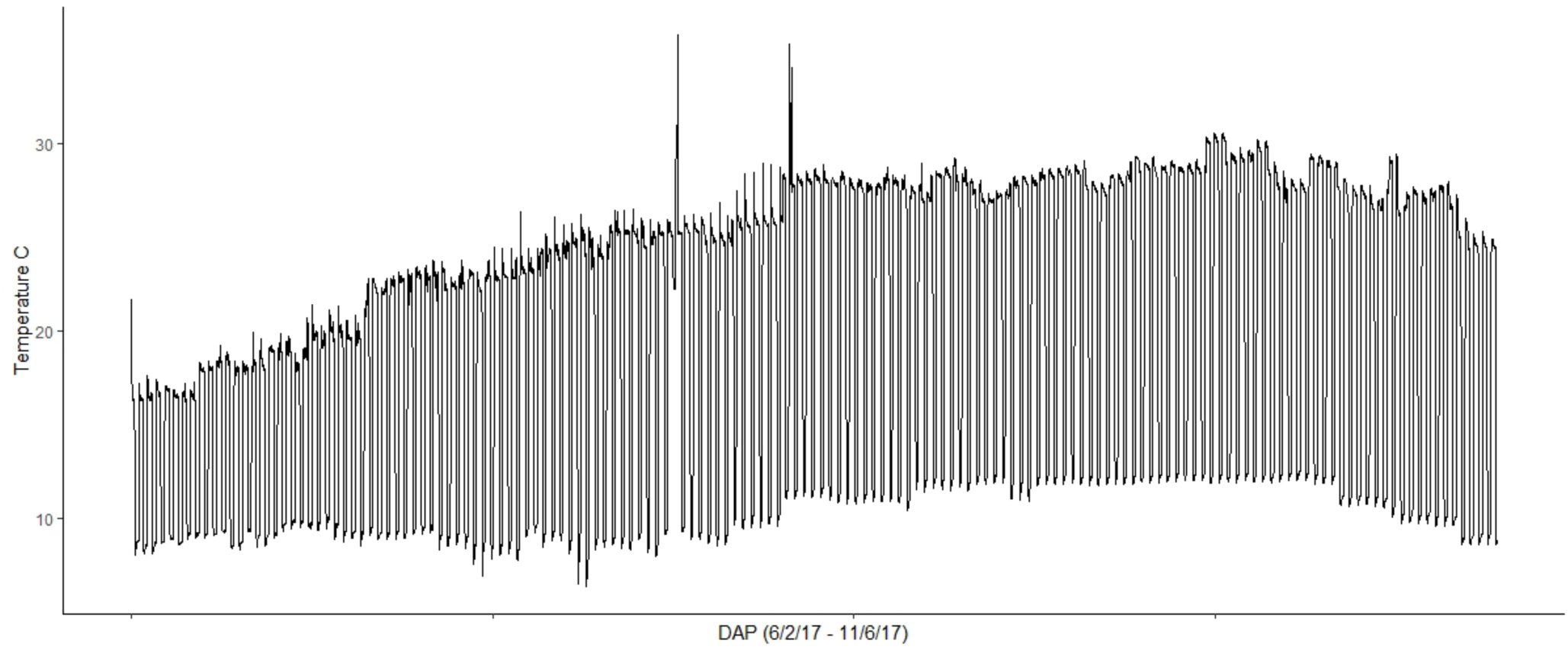
Time series of temperature (°C) conditions using a HOBO data logger (HOBO®, Onset Computer Corporation). Temporary spikes in temperature were due to brief power outages and did not result in any clear detrimental effects on experimental plants.

**Figure S4.**
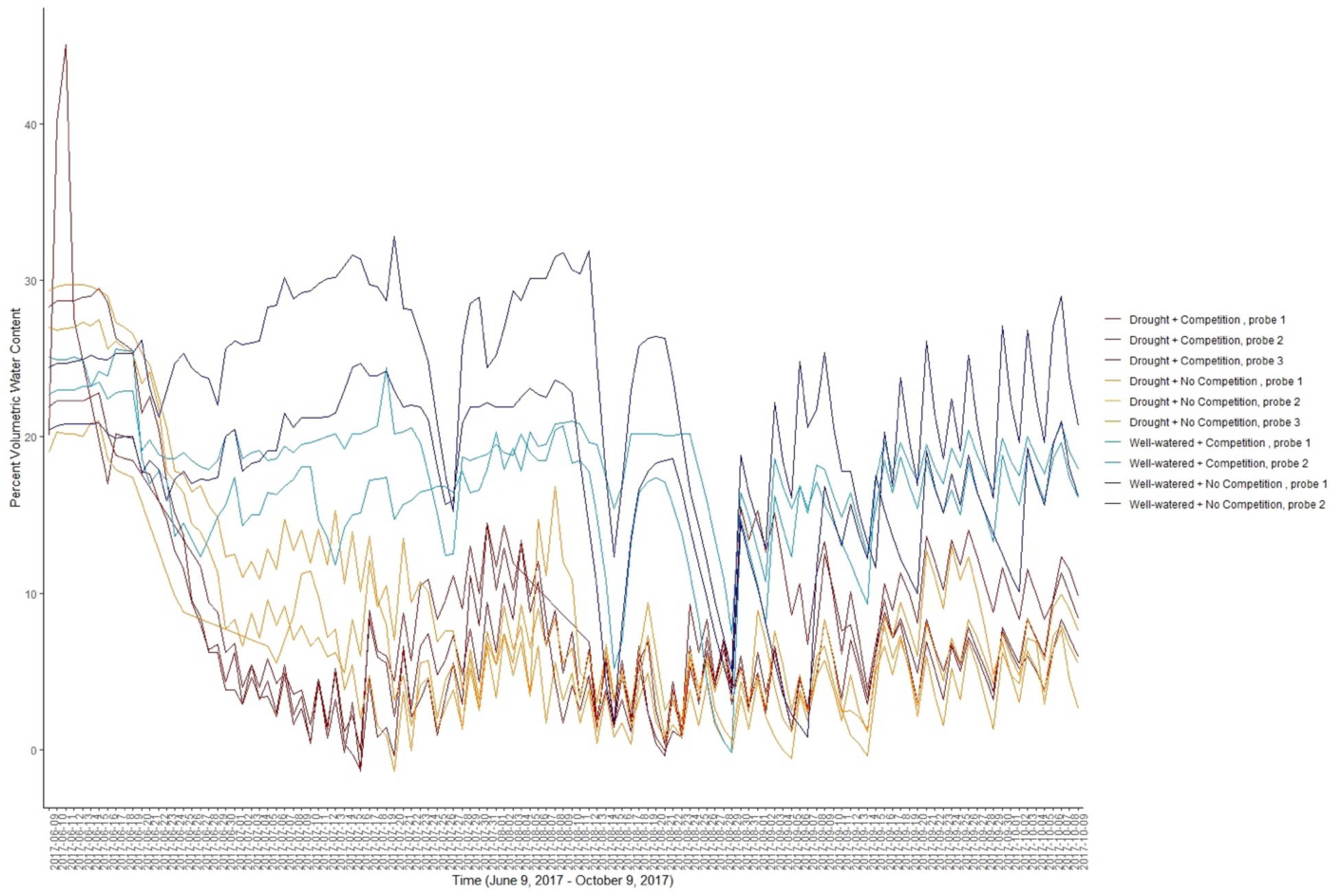
Percent volumetric water content (%VWC) of potting mix within pots for each treatment was monitored using EM-50 dataloggers and EC-5 probes (METER Group Inc., Pullman, WA). Seeds were planted on May 31, %VWC logging was started on June 9, and drought treatments were started on June 20.

**Figure S5.**
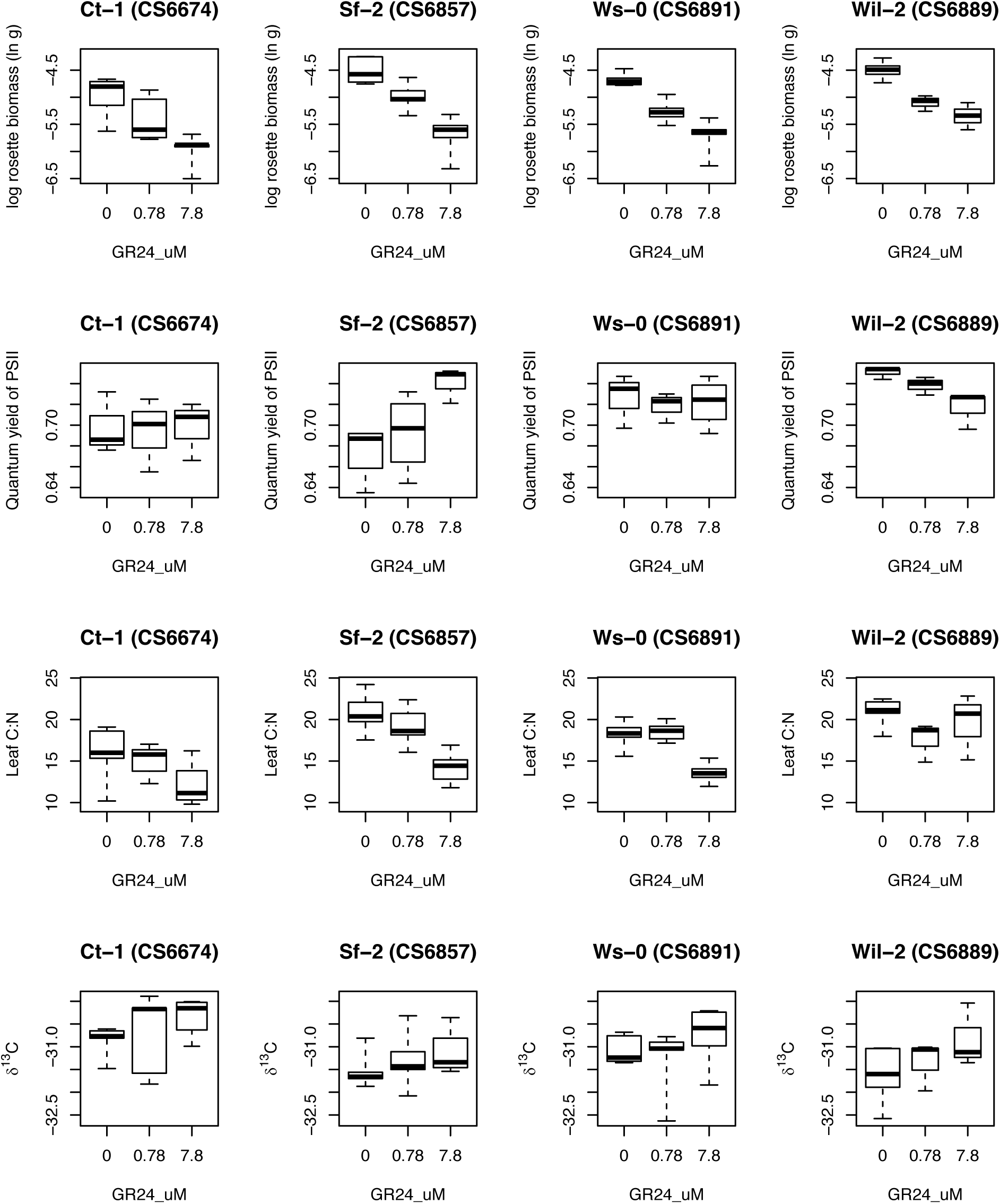
Response of four genotypes to increasing concentrations of GR24 applied to soil.

**Figure S6.**
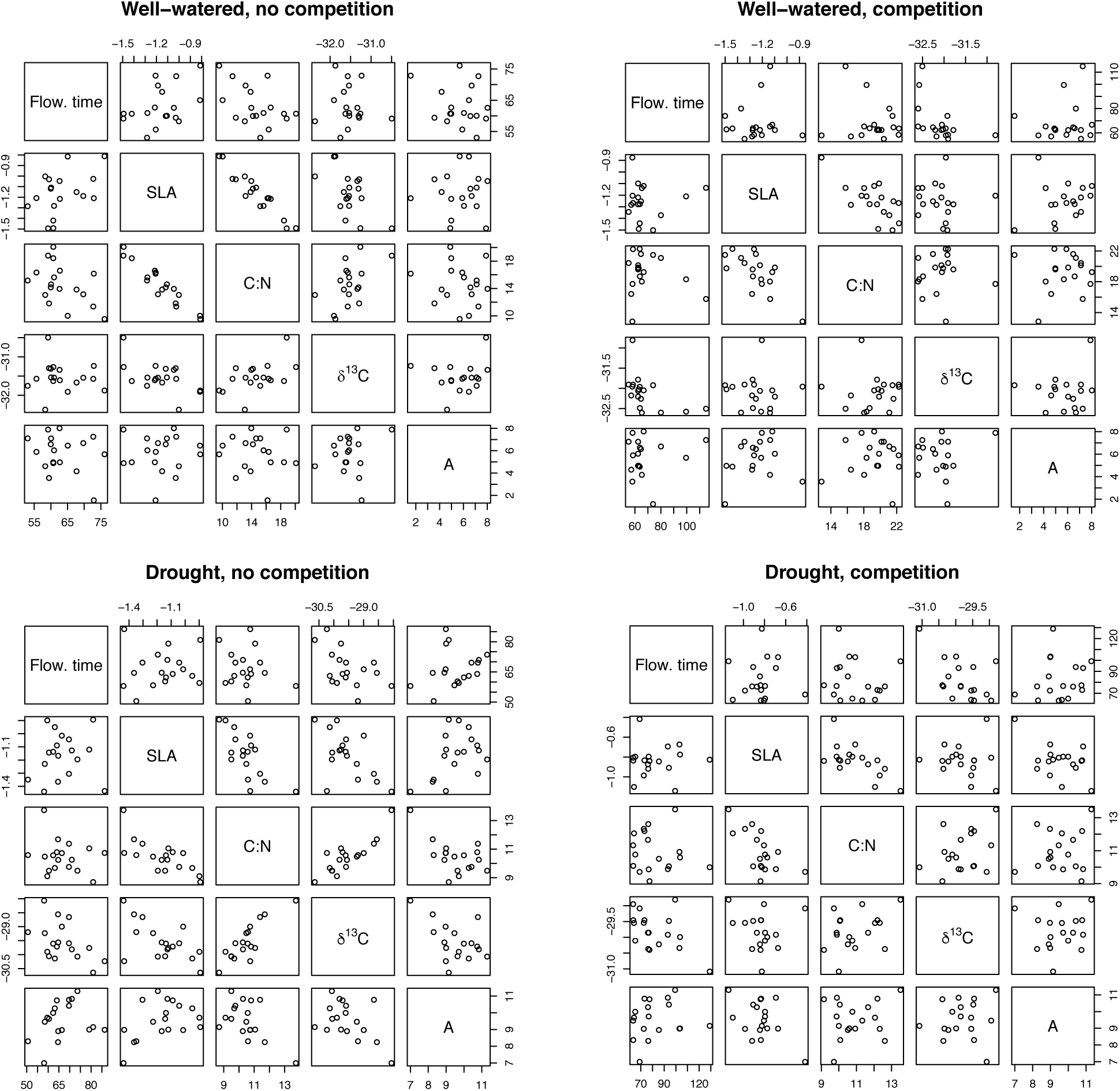
Bivariate relationships between traits often thought to be exhibit coordinated variation. Note that gas exchange (used to caculate *A*) was only measured in the absence of competitors, these estimates are used in the comparisons with traits under competition.

**Figure S7.**
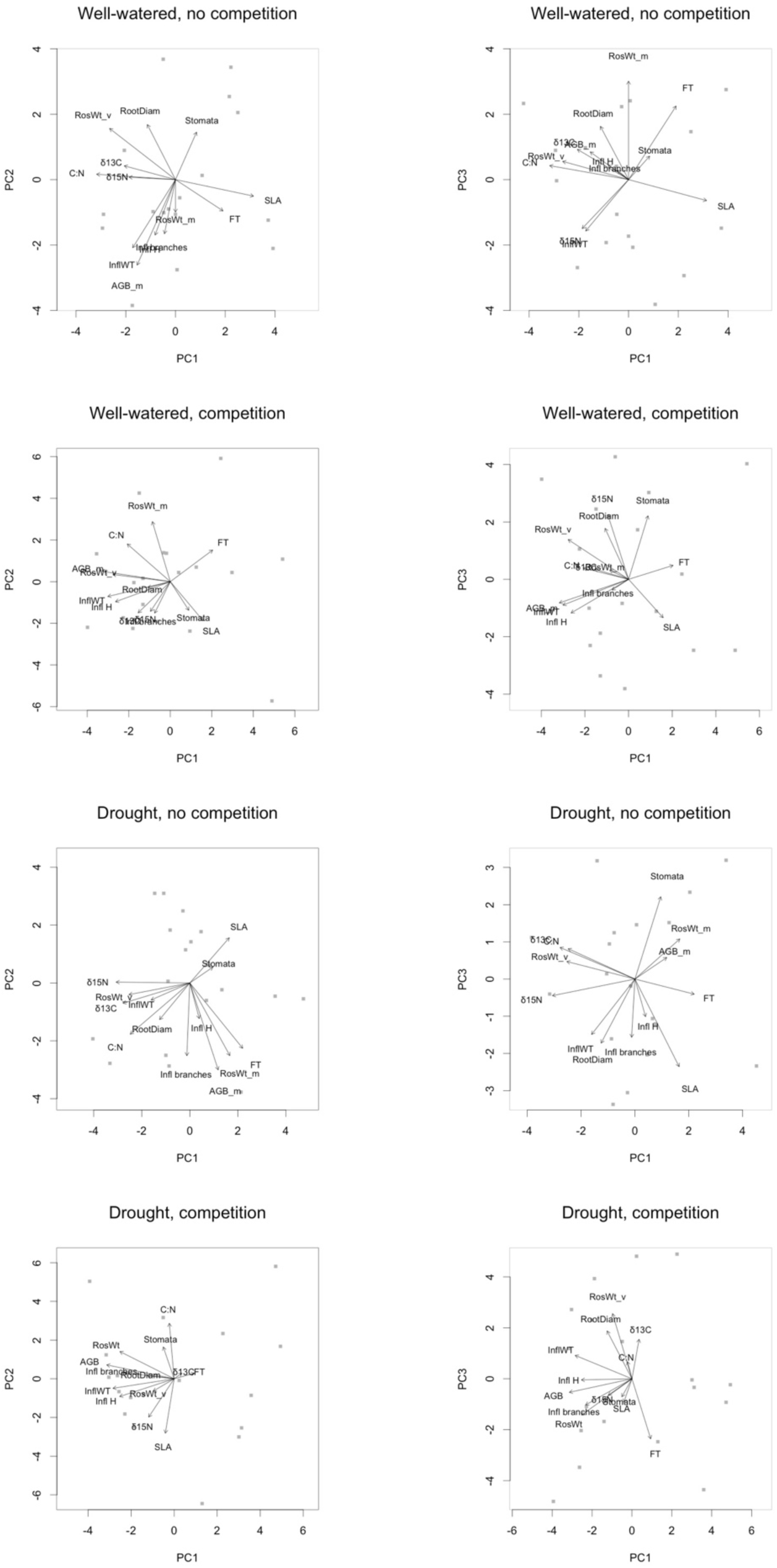
Principal components of trait variation within each treatment.

**Table S1.**
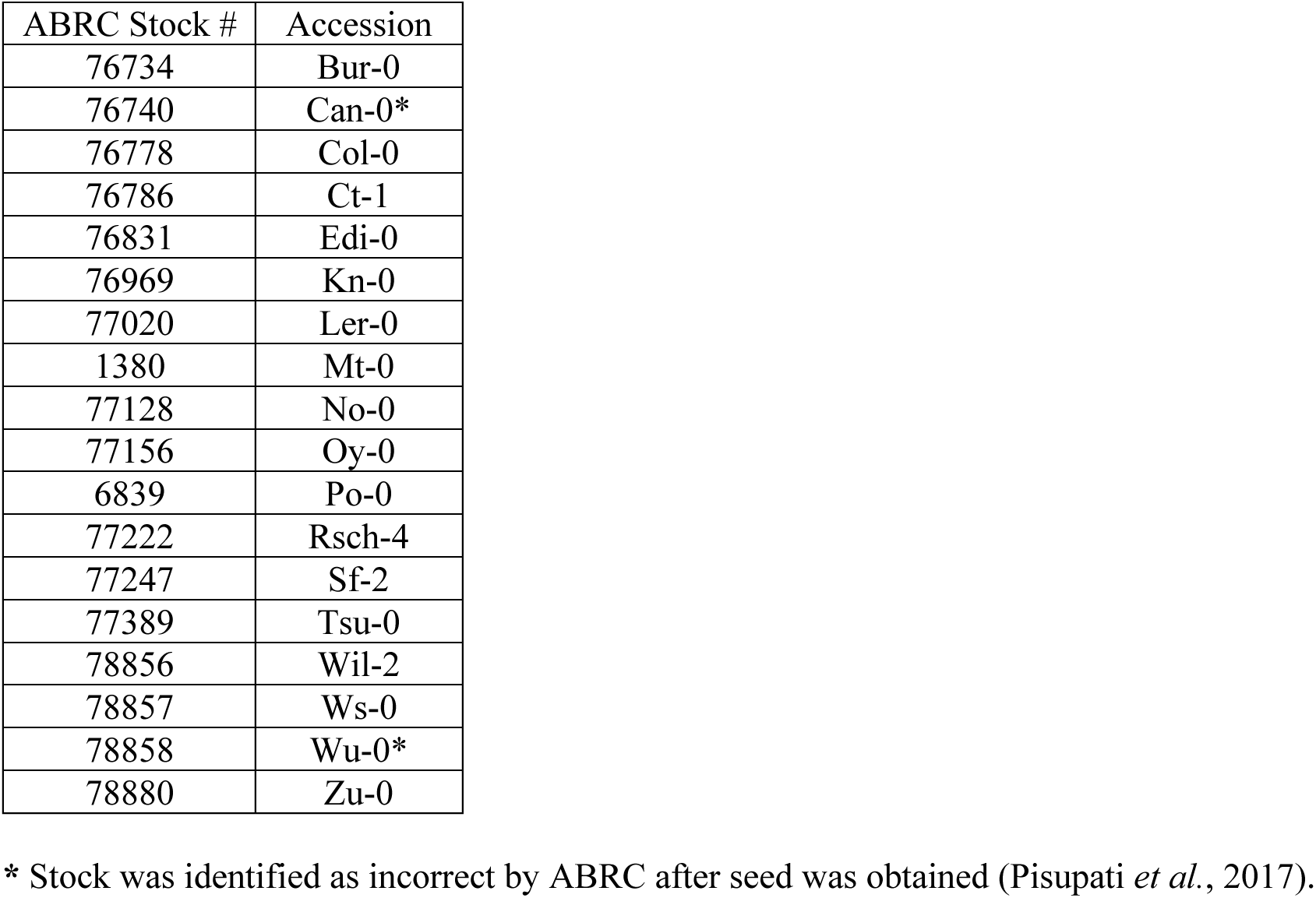
The 18 parent genotypes from the Multiparent Advanced Generation Inter-Cross (MAGIC) used in this study.

**Table S2.** Climate mean conditions for non-contaminant ecotypes using WorldClim (Hijmans et al. 2005) and CGIAR CSI Aridity (Zomer et al. 2008) data.

| Line | Longitude | Latitude | Alt | Ann.Mean.Tmp<br>(°C) | Ann.Prc<br>(mm) | Ann.Prc/PET |
| --- | --- | --- | --- | --- | --- | --- |
| Bur-0 | -6.2 | 54.1 | 172 | 8.4 | 1068 | 1.8098 |
| Can-0* |  |  |  |  |  |  |
| Col-0 |  |  |  |  |  |  |
| Ct-1 | 15 | 37.3 | 40 | 17.5 | 503 | 0.4209 |
| Edi-0 | -3.16028 | 55.9494 | 149 | 8.3 | 743 | 1.2129 |
| Hi-0 | 5 | 52 | -2 | 9.4 | 800 | 1.173 |
| Kn-0 | 23.8924 | 54.8969 | 30 | 6.7 | 613 | 0.9068 |
| Ler-0 | 10.8719 | 47.984 | 621 | 8 | 962 | 1.2847 |
| Mt-0 | 22.46 | 32.34 | 282 | 17.5 | 132 | 0.1044 |
| No-0 | 13.2995 | 51.0581 | 276 | 8.1 | 627 | 0.8947 |
| Oy-0 | 6.193019 | 60.385543 | 5 | 7.2 | 2190 | 4.4512 |
| Po-0 | 7.09 | 50.72 | 67 | 10 | 739 | 0.97 |
| Rsch-4 | 34 | 56.3 | 210 | 4.1 | 640 | 1.0126 |
| Sf-2 | 3.03333 | 41.7833 | 146 | 15.4 | 677 | 0.765 |
| Tsu-0 | 136.31 | 34.43 | 160 | 14.9 | 2385 | 2.319 |
| Wil-2 | 25.3167 | 54.6833 | 106 | 6.4 | 656 | 0.988 |
| Ws-0 | 30 | 52.3 | 130 | 6.7 | 624 | 0.8398 |
| Wu-0* |  |  |  |  |  |  |
| Zu-0 | 8.55 | 47.3667 | 424 | 9.3 | 1087 | 1.4135 |

**Table S3.** Temperature and photoperiod program for Conviron growth chamber, to simulate early spring germination for a spring annual.

| Week | Night °C | Day °C | Photoperiod (Hours:Minutes) |
| --- | --- | --- | --- |
| 1 | 10 | 15 | 13:04 |
| 2 | 10 | 15 | 13:30 |
| 3 | 10 | 15 | 13:54 |
| 4 | 10 | 16 | 14:20 |
| 5 | 10 | 17 | 14:42 |
| 6 | 10 | 18 | 15:06 |
| 7 | 10 | 19 | 15:26 |
| 8 | 10 | 20 | 15:44 |
| 9 | 10 | 21 | 16:00 |
| 10 | 10 | 21 | 16:12 |
| 11 | 11 | 22 | 16:18 |
| 12 | 12 | 23 | 16:22 |
| 13 | 12 | 23 | 16:20 |
| 14 | 13 | 24 | 16:16 |
| 15 | 13 | 24 | 16:06 |
| 16 | 13 | 24 | 15:52 |
| 17 | 13 | 24 | 15:34 |
| 18 | 13 | 24 | 15:16 |
| 19 | 13 | 24 | 14:54 |
| 20 | 13 | 24 | 14:32 |
| 21 | 12 | 23 | 14:08 |
| 22 | 11 | 22 | 13:42 |
| 23 | 10 | 21 | 13:18 |
| 24 | 10 | 20 | 12:52 |

**Table S4.** Broad sense heritability of traits within each treatment. ‘m’ suffixes indicate mature end stage harvest, while ‘v’ indicates the pre-flowering vegetative stage harvest.

|  | WW NC | WW C | D NC | D C |
| --- | --- | --- | --- | --- |
| Absolute fitness | 0.7880 | 0.7337 | 0.5693 | 0.2797 |
| AGB_m | 0.6026 | 0.5489 | 0.6739 | 0.4255 |
| Rosette wt_m | 0.8592 | 0.5178 | 0.7970 | 0.4030 |
| Inflorescence wt | 0.6081 | 0.6419 | 0.3673 | 0.4387 |
| Flowering time | 0.8932 | 0.9568 | 0.8873 | 0.8798 |
| Inflorescence height | 0.8378 | 0.7176 | 0.4076 | 0.3531 |
| Inflorescence branches | 0.6920 | 0.4795 | 0.5927 | 0.3241 |
| Siliques number | 0.7972 | 0.6835 | 0.5705 | 0.3024 |
| Average siliques length | 0.5640 | 0.6601 | 0.4642 | 0.2772 |
| Root wt_v | 0.3012 |  | 0.1627 |  |
| Root diam_v | 0.2952 | 0.2444 | 0.1254 | 0.4691 |
| Root:shoot | 0.3087 |  | 0.2019 |  |
| Total leaf area_v | 0.3101 | 0.3695 | 0.2286 | 0.3768 |
| Stomatal density | 0.2026 | 0.2080 | 0.3614 | 0.3554 |
| Rosette wt_v | 0.6003 | 0.4206 | 0.2611 | 0.3754 |
| log(SLA) | 0.3924 | 0.2513 | 0.3884 | 0.2936 |
| Leaf prop N | 0.5285 | 0.3096 | 0.2566 | 0.3425 |
| Leaf prop C | 0.2960 | 0.2139 | 0.2566 | 0.2951 |
| Leaf C:N | 0.5785 | 0.4204 | 0.3269 | 0.3335 |
| $\delta^{13}\text{C}$ | 0.3825 | 0.4691 | 0.3838 | 0.2766 |
| $\delta^{15}\text{N}$ | 0.2820 | 0.2496 | 0.3640 | 0.3437 |
| *A* (whole plant assimilation) | 0.1325 |  | 0.1633 |  |
| Brachypodium AGB |  | 0.2780 |  | 0.3606 |

Table S5. Variance components estimates for linear mixed-effects model where genotype, genotype-treatment interactions, and tray effects were modeled as random effects and treatments were fixed effects. ‘est’ refers to the variance component estimate and ‘LCL’ and ‘UCL’ refer to the lower and upper 95% confidence limits for the estimate. Variance components estimated and fixed at zero report ‘NA’ for the 95% CI.

See attached files.

**Table S6.** Tests for changing selection between competition treatments within the same moisture treatment. Slopes give the relationship between traits and relative fitness, where relative fitness is standardized **across** the treatment combination of competition and moisture conditions (corresponding to a model of hard selection), and traits are standardized across competition treatments being compared. Bold indicates those where slopes differ significantly between competition treatments with FDR < 0.05. For results when fitness is relativized across environments see Table 3.

| *Trait* | *Well-watered* | *Slope, w/ comp* | *Slope, no comp.* | *t-test p* | *Drought* | *Slope, w/ comp* | *Slope, no comp.* | *t-test p* |
| --- | --- | --- | --- | --- | --- | --- | --- | --- |
| AGB_m |  | 0.94 | 0.24 | 0.3812 |  | 0.25 | -0.97 | 0.2369 |
| Rosette wt_m |  | -0.28 | -0.53 | 0.6964 |  | -0.15 | -0.73 | 0.5193 |
| Inflorescence wt |  | 0.81 | 0.79 | 0.9597 |  | 0.48 | 0.80 | 0.6863 |
| Inflorescence height |  | 0.18 | 0.02 | 0.4455 |  | -0.01 | -0.09 | 0.7722 |
| Inflorescence branches |  | 0.23 | -0.07 | 0.3425 |  | 0.02 | -0.38 | 0.3158 |
| Flowering time |  | -0.08 | -0.86 | **<0.0001** |  | -0.08 | -0.68 | 0.0060 |
| Silique number |  | 0.78 | 0.73 | 0.6411 |  | 0.68 | 0.83 | 0.3220 |
| Average silique length |  | 0.08 | 0.04 | 0.7673 |  | 0.13 | 0.53 | 0.0810 |
| Root diam_v |  | 0.06 | -0.10 | 0.5152 |  | 0.06 | -0.26 | 0.5425 |
| Stomatal density |  | -0.05 | -0.21 | 0.2436 |  | -0.08 | 0.06 | 0.4271 |
| Rosette wt_v |  | 0.39 | 0.26 | 0.8172 |  | 0.38 | -0.42 | 0.5942 |
| log(SLA) |  | -0.01 | -0.15 | 0.3117 |  | -0.01 | 0.23 | 0.3230 |
| A (whole plant assimilation) |  | 0.04 | 0.12 | 0.5418 |  | 0.00 | -0.12 | 0.5139 |
| Leaf C:N |  | 0.06 | 0.25 | 0.2721 |  | 0.01 | -0.08 | 0.6113 |
| δ13C |  | 0.07 | 0.09 | 0.9331 |  | 0.09 | 0.07 | 0.9008 |
| δ15N |  | 0.02 | 0.34 | 0.4730 |  | 0.03 | 0.12 | 0.6353 |

**Table S7.** Selection on mean and plasticity of trait values across competition treatments. Plasticity was calculated as no competition – competition treatments. For example, selection for an increase in inflorescence height in well-watered competition treatments is indicated by a negative coefficient. None of these tests for selection on plasticity were significant using a false discovery rate of 0.05 within each water treatment.

| *Trait* | **Well-watered** |  |  |  | **Drought** |  |  |  |
| --- | --- | --- | --- | --- | --- | --- | --- | --- |
|  | *Trait*<br>*avg.*<br>*coeff.* | *Trait*<br>*avg. p* | *Trait*<br>*plast.*<br>*coeff.* | *Trait*<br>*plast. p* | *Trait*<br>*avg.*<br>*coeff.* | *Trait*<br>*avg. p* | *Trait*<br>*plast.*<br>*coeff.* | *Trait*<br>*plast. p* |
| AGB_m | 0.2650 | 0.0308 | -0.2166 | 0.0705 | -0.3588 | 0.0209 | 0.1413 | 0.3254 |
| Rosette wt_m | -0.1773 | 0.4933 | -0.0116 | 0.9638 | -0.6082 | 0.0057 | 0.3485 | 0.0848 |
| Inflorescence<br>wt | 0.3517 | 0.0019 | -0.1323 | 0.1775 | -0.0133 | 0.9222 | 0.2081 | 0.1414 |
| Flowering time | -0.3877 | 0.0004 | -0.1145 | 0.1950 | -0.4004 | 0.0105 | -0.2510 | 0.0870 |
| Inflorescence<br>height | 0.1203 | 0.0602 | -0.2056 | 0.0034 | -0.0746 | 0.4506 | 0.0756 | 0.4450 |
| Inflorescence<br>branches | 0.0818 | 0.4116 | -0.1058 | 0.2915 | -0.0956 | 0.4823 | -0.0801 | 0.5552 |
| Silique number | 0.3852 | 0.0000 | -0.0667 | 0.1681 | 0.3362 | 0.0002 | 0.0323 | 0.6357 |
| Average<br>silique length | 0.1113 | 0.1363 | -0.1480 | 0.0537 | 0.2355 | 0.0135 | -0.0003 | 0.9970 |
| Root diam_v | 0.0491 | 0.5826 | -0.1139 | 0.2125 | -0.0649 | 0.5133 | 0.0386 | 0.6958 |
| Stomatal<br>density | -0.1108 | 0.1869 | -0.0565 | 0.4912 | -0.0602 | 0.5424 | 0.1225 | 0.2241 |
| Rosette wt_v | 0.1423 | 0.5122 | -0.0109 | 0.9597 | 0.0963 | 0.6021 | -0.1698 | 0.3622 |
| log(SLA) | -0.0364 | 0.6589 | -0.1200 | 0.1582 | 0.0799 | 0.4150 | 0.0448 | 0.6451 |
| Leaf C:N | 0.0995 | 0.2216 | 0.0905 | 0.2638 | -0.0327 | 0.7434 | 0.0112 | 0.9108 |
| $\delta^{13}\text{C}$ | 0.0964 | 0.2224 | -0.1133 | 0.1553 | 0.1617 | 0.0817 | -0.1151 | 0.2039 |
| $\delta^{15}\text{N}$ | 0.0346 | 0.6815 | 0.0762 | 0.3714 | 0.0688 | 0.4971 | 0.0356 | 0.7241 |

